# Common Dysregulated Pathways and Genes in Alzheimer’s, Parkinson’s, and Huntington’s Diseases: A Computational Analysis to Nominate Candidate Gene Targets for Drug Repurposing

**DOI:** 10.64898/2026.07.22.739294

**Authors:** Nehal Adel Abdelsalam, Muhammad Elsadany, Eman Badr

## Abstract

Neurodegenerative diseases are a major threat to older adults and represent a growing global health burden as the elderly population continues to expand. The complexity of neurodegenerative diseases and the incomplete understanding of their pathophysiology limit the development of effective therapeutics. To study common neurodegenerative mechanisms and accordingly potential drug targets across Alzheimer’s, Parkinson’s, and Huntington’s diseases, transcriptomic profiles of patients with each disease were analyzed to detect common differentially expressed genes and common enriched pathways. Common differentially expressed genes involved in the shared pathways were identified as key drug targets and validated in silico to study the impact of their dysregulation. Three pathways were enriched and upregulated across the three diseases, along with 274 common differentially expressed genes. Two of the shared pathways were involved in activation of the transcription factor nuclear factor kappa B (NF-κB), indicating inflammatory signaling. The third pathway involved regulation of BCL2L11 transcription by RUNX3, which contributes to the protective effect of neurodegenerative diseases against cancer. Five key drug targets were identified: *NFKBIA*, *NFKB1*, *RELA*, *TRIM4*, and *SMAD4*. They were significantly upregulated across all three diseases and involved in the shared pathways. Drugs that target the expression of these genes and previously approved by Food and Drug Administration were reported for treatment repurposing for the three neurodegenerative diseases. The resultant drug list included the conventional and commonly safe antihyperlipidemics, antihypertensives, antidiabetics, analgesics, diuretics, antiparkinson’s, and antipsychotics.

## Introduction

Neurological disorders cause 16.5% of deaths and 11.6% of disability-adjusted life years (1). Neurodegeneration is a neurological disorder in which a gradual loss of the structure or function of the nerve cells occurs due to cell death, axonal regeneration failure, or neuron demyelination. These nerve disorders are generally referred to as neurodegenerative diseases (NDs). NDs are either partial or complete, solo or combined, hereditary or acquired, known or unknown in origin (2). They are a global burden that mainly threatens the elderly (3). In addition to aging, diverse pathways involved in intracellular processes, the local tissue environment, and the systemic environment can play a pivotal role in developing the NDs (4). Mitochondrial malfunction, cellular stress, dysfunctional cytoskeletal proteins, dysregulated programmed cell death, and changes in cellular protein homeostasis are all probable drivers for the imbalance between neuronal survival and degeneration (5–13). NDs affect different brain areas and manifest distinctly common features, such as a gradual loss of sensory, motor, or cognitive abilities (14). Research on the common mechanisms behind different NDs expands the horizon for new or repurposed therapeutic agents that may simultaneously treat multiple diseases, for example: Alzheimer’s (AD), Parkinson’s (PD), and Huntington’s diseases (HD). These are primarily classified as proteinopathies NDs as they are associated with the aggregation of misfolded proteins (15).

Alzheimer’s disease (AD) starts as a mild memory loss, with symptoms worsening over time. The buildup of extracellular β-amyloid protein and intracellular tau protein is one of the main reasons behind AD (16). A recent bioinformatic study identified 1031 unique, differentially expressed genes (DEGs) that implicate all major cell types in AD patients compared to healthy controls. While DEGs were downregulated by 75% in excitatory neurons and 95% in inhibitory neurons, most of the identified DEGs in astrocytes, oligodendrocytes, and microglia were upregulated within a range of 53%–63% (17).

Parkinson’s disease (PD) is a movement disorder characterized by aggregations of α-synuclein that form Lewy bodies and neurites (18). The disease hallmark symptom of the disease is tremors in hands, arms, legs, and neck muscles. Studies showed that the increase of the wild type of *SNCA* gene copy number induced elevated levels of α-synuclein expression and could contribute to developing PD (19,20). In another study, 85 genes were identified as significantly hypo-methylated and upregulated in patients with PD compared to healthy controls. Downregulated genes were significantly associated with the structural constituent of the cytoskeleton and, thus, dopaminergic neurotransmission. Upregulated genes were associated with phagosome and lysosome pathways and involved in misfolded protein degradation (21).

Huntington’s disease (HD) is a neurodegenerative disease that causes chorea, cognitive impairment, physical and mental disturbances, and probable depression (22). It is induced by the intracellular accumulation of mutant Huntington’s protein aggregates, resulting in brain cell apoptosis, mainly in the striatum (22). On the genetic level, HD is an autosomal dominant neurodegenerative disease caused by a CAG repeat expansion in exon 1 of the Huntingtin gene, which results in a polyglutamine strand at the Huntingtin protein’s N-terminus (23). In a study, a bioinformatics analysis on HD was performed, and sixty genes were specified as differentially expressed in HD gene carriers compared with healthy controls. A total of 109 enriched pathways were identified using the Gene Ontology (GO) analysis of the list of DEGs and *ADGRG1* gene was among the top genes involved in the enriched pathways (24). These pathways included vascular endothelial growth factor production, cerebral cortex radial glia-guided migration, layer formation in the cerebral cortex, and cerebral cortex regionalization. In addition, different GO terms involved in neurodevelopment were identified, such as the positive regulation of neuron migration cell adhesion (24).

NDs are considered incurable as there is no known intervention to stop the gradual destruction of neurons and full pathophysiological mechanisms linked to NDs are still unknown (25). Meanwhile, the known underlying mechanisms are polyfactorial and result from the complicated interactions of the innumerable partially unknown genetic and non-genetic factors (25). Although these diseases have distinct clinical manifestations, they all diminish brain functionality due to irreversible damage in neurons (26). Previous research investigated the common proteins and pathways between two or more NDs (27,28). Neuroinflammation and oxidative stress were found to be strong, common candidates for neurodegeneration (29). The identified common proteins and pathways were proposed as potential drug targets for treatment (28). It is worth mentioning that multi-target drugs for NDs are a promising strategy for treatment, offering higher efficacy, fewer side effects, and reduced resistance to treatment (30). Hence, in this study, we aimed to identify common pathways and drug targets among the three neurodegenerative diseases under study. We conducted a bioinformatics pipeline for pathway enrichment and differential gene expression to study Alzheimer’s, Parkinson’s, and Huntington’s diseases, as summarized in Fig. 1.

**Fig. 1.**
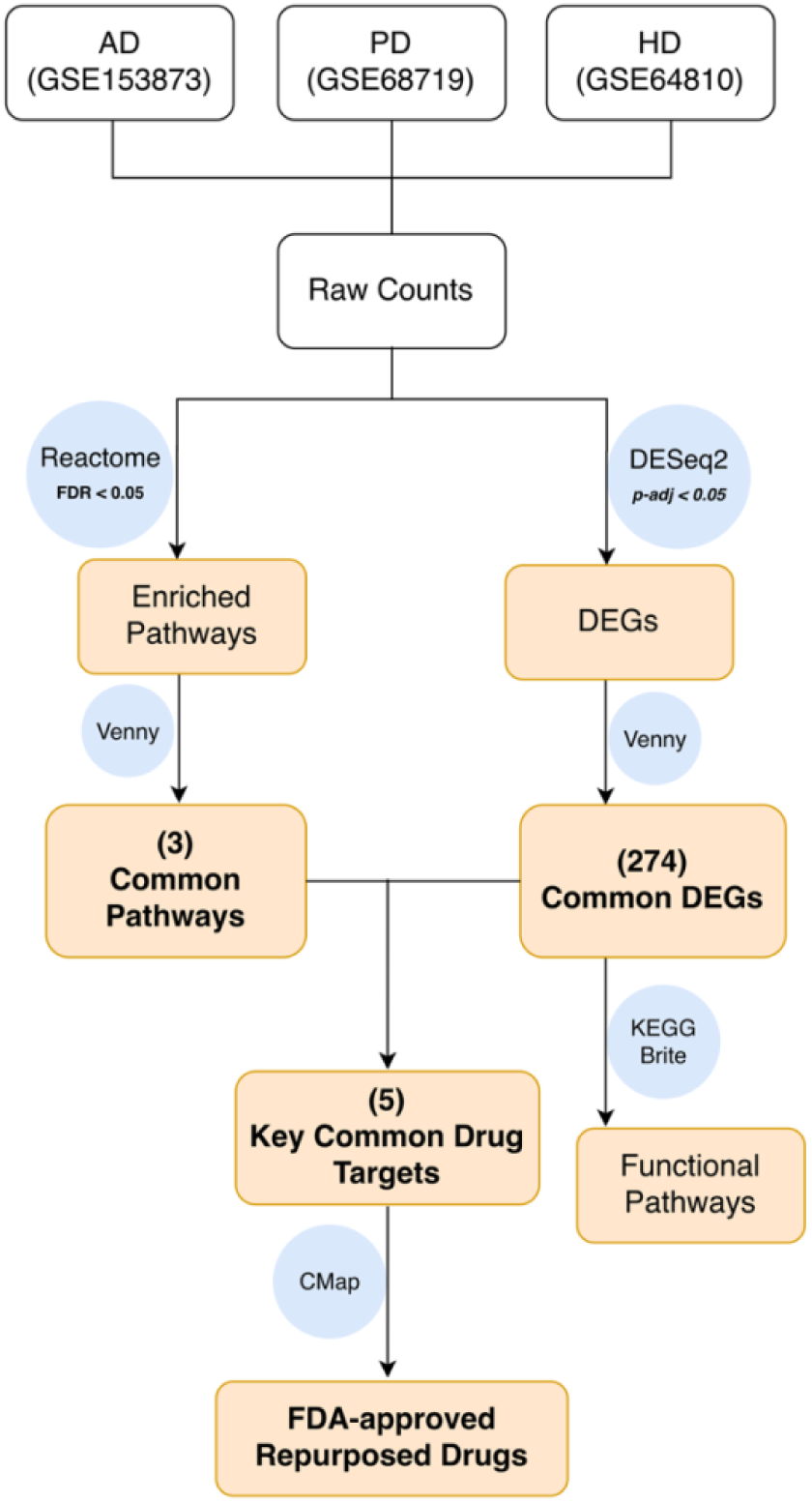
**Analysis pipeline overview (FDR: False discovery rate; p-adj: Adjusted P-value; DEGs: Differentially expressed genes; FDA: Food and Drug Administration).**

We report the mutually enriched pathways across the three diseases and shed light on the common pathological processes to help repurpose drugs for a broader range of NDs, rather than investigating each disease independently. Three mutually enriched pathways were reported based on the raw expression counts of each disease. Two of these pathways were involved in activating the transcription factor, activated nuclear factor kappa B (NF-κB). The third pathway was the regulation of BCL2L11 (BIM) transcription by RUNX3, which was found to correlate with cancer progression. We conducted differential expression analysis using raw counts to conclude mutual DEGs that were involved in the identified mutual pathways. This set of mutual DEGs was defined as “key mutual drug targets,” and they were internally and externally validated in silico for impact on the three disease states. Food and Drug Administration (FDA)-approved drugs targeting the expression of this gene set were sought to be repurposed for the treatment of the three diseases, employing a multi-target strategy.

## Methods

### Datasets description

Gene Expression Omnibus (GEO) database (https://www.ncbi.nlm.nih.gov/geo/) was searched for genome-wide expression datasets (31). We aimed to include datasets with comparable sample sizes and experimental design where raw expression counts were available. The search strategy was as follows: <name of disease> AND “RNA-seq” AND “human”, where the name of the disease is either “Alzheimer’s”, “Parkinson’s”, or “Huntington”. The dataset inclusion criteria were: (a) all datasets were genome-wide, (b) the GEO series type was expression profiling by high throughput sequencing, (c) raw data files were available, with control and disease samples of the human brain, (d) all samples were tissue samples and not blood-derived. Hence, the three retrieved datasets were: GSE153873 for AD, GSE68719 for PD, and GSE64810 for HD.

The AD dataset (GSE153873) had three groups: the AD group, the old control group, and the young control group. The AD group consisted of 12 samples. The healthy controls were divided into the old control group (10 samples) and the young control group (8 samples). The age of young controls ranged from 42 years to 60 years, while the old control and AD groups ranged from 61-79 years. The brain tissue samples were obtained from postmortem human brain samples from the lateral temporal lobe (Brodmann area 21 or 20) (32). The AD and old age control were included in the analysis to compare patients with AD to healthy control individuals at similar age.

The PD dataset (GSE68719) was divided into the PD and neurologically normal control groups. The PD group comprised 29 samples, and the healthy control group included 44 samples. The age of healthy controls ranged from 46 to 97 years, while the patients with PD ranged from 64 to 95 years. The brain tissue samples were obtained from postmortem human brain samples from the prefrontal cortex Brodmann area 9 (33).

The HD dataset (GSE64810) had two groups: the HD group (20 samples) and the healthy control group (49 samples). The age of the healthy control group ranged from 36 to 106 years, while the HD patients’ age ranged from 40 to 75 years. The brain tissue samples were obtained from postmortem human brain samples from the prefrontal cortex Brodmann area 9 (34).

### Pathway enrichment analysis

Pathway enrichment analysis was conducted on the RNA-Seq raw counts for each disease. A pathway was considered enriched with a false discovery rate (FDR) below 0.05 using the Reactome Pathway Browser Tool (https://reactome.org/PathwayBrowser/#TOOL=AT)(35), where the overall log fold change (lfc) for all genes involved in a pathway is calculated. The Venny online tool (https://bioinfogp.cnb.csic.es/tools/venny/index.html) was used to identify the mutual pathways between the three diseases (36). Three distinct pathways were found to be mutual between the NDs under study.

### Differential gene expression analysis and functional pathway analysis

The differential gene expression analysis was performed using DESeq2 R package v.1.30.0 (37). Genes with adjusted P-value less than 0.05 were defined to be significantly differentially expressed. After matching the DEGs with their Ensembl IDs, the org.Hs.eg.db R package (version 3.12.0) was used to convert Ensembl IDs to gene symbols (38). This step was performed in two datasets, PD and HD because gene symbols were already provided in the AD dataset. Volcano plots visualized gene expression in each disease using the EnhancedVolcano R package (https://github.com/kevinblighe/EnhancedVolcano) after applying log fold change shrinkage on counts using ashr (39). To conclude the mutual DEGs between the three NDs, the Venny online tool (36) was used, and a Venn diagram was plotted.

Kyoto Encyclopedia of Genes and Genomes (KEGG) Brite database was used to get functional classification systems for the mutual DEGs between the three diseases (40). The color tool was used to access the KEGG Brite database to navigate the mutual DEGs against KEGG pathway maps, KEGG Brite hierarchies, and KEGG modules.

### Exploration and validation of the key mutual drug targets between the three diseases

We sought genes that were key players in the three diseases to serve as potential drug targets, defined as “key mutual drug target”. To be considered a key mutual drug target, a gene had to be involved in a mutual enriched pathway and differentially expressed with the same pattern across the three diseases. The identified genes were tested for their diagnostic potential by the receiver operator characteristics (ROC) curve analyses. ROC curves were visualized using the pROC R package (41). As an internal validation, ROC analysis was performed using the study datasets. For further in-silico validation, these genes were tested by ROC using the external datasets GSE184942, GSE106608, and GSE97100 for AD, PD, and HD, respectively.

### Drug repurposing for the key mutual targets

To identify the potential drugs that could reverse the expression of the key mutual drug targets, Connectivity Map (CMap) (https://www.broadinstitute.org/connectivity-map-cmap) data from the broad institute were used (GSE70138) (42). The CMap dataset provides the transcriptomic profiles of genes in response to different perturbagens (e.g., drugs, chemical compounds, and reagents) after being tested on different human cell lines. This dataset was filtered to include only FDA-approved drugs and the key mutual drug targets. The drugs were then filtered to retain only those that simultaneously reversed the expression of the key mutual drug targets. We further filtered the resultant drugs to exclude discontinued drugs, topical medications, chemotherapeutics, antibiotics, antifungals, antivirals, hormone therapies, anesthetics, antiplatelet agents, anticoagulants, and medications used for rare diseases.

## Results

### Biological pathways enriched uniquely in each ND: inflammation, apoptosis, and neurotransmission

Pathway enrichment analyses were conducted on the raw RNA-seq counts for each neurodegenerative disease using Reactome and an FDR threshold less than 0.05. Table 1 summarizes the number of significantly enriched pathways. A full list of the pathways is present in the Supplementary Information tables S1.1, S1.2, and S1.3 (Additional File 1).

**Table 1:** Summary of the enriched pathways in each ND.

| Enriched Pathways (FDR<0.05) |  |  |  |
| --- | --- | --- | --- |
|  | Upregulated | Downregulated | Total |
| AD | 69 | 84 | 153 |
| PD | 76 | 39 | 115 |
| HD | 80 | 29 | 109 |

The AD enriched pathways were investigated to highlight the dysregulated and impaired pathways for patients with AD. The pathway R-HSA-9022702 (MECP2 regulates the transcription of neuronal ligands) was found to be downregulated with an FDR value of 0.001. *MECP2* regulates the transcription of several transcription factors involved in the functioning of the nervous system, such as *CREB1* and *MEF2C*. It was shown that increased dosage or loss of function of the *MECP2* gene could cause a plethora of neuropsychiatric disorders (43). It was demonstrated that *MECP2* could increase the pro-inflammatory response of microglial cells. In postmortem brain samples from different stages of AD, it was found that phosphorylation of the *MECP2* protein decreased at the amino acid serine 423 in the early stages of Alzheimer’s disease (44). Another study identified MECP2 as a possible regulator of Tau pathology in mouse models (45).

The enrichment analysis showed that the pathway R-HSA-264642, the acetylcholine neurotransmitter release cycle, was downregulated. This cycle involves the synthesis of acetylcholine, loading of synaptic vesicles, docking and priming of the acetylcholine-loaded synaptic vesicles, and then release of acetylcholine (46). Other pathways involved in neurotransmitters secretion and regulation, such as R-HSA-888590 (GABA synthesis, release, reuptake, and degradation) and R-HSA-181429 (serotonin neurotransmitter release cycle), were also downregulated with significant FDR values.

Pathways involved in apoptosis were significantly upregulated in the transcriptome analysis of patients with AD. Examples are R-HSA-111458 (formation of apoptosome) and R-HSA-9627069 (Regulation of the apoptosome activity). R-HSA-205017 (NFG and proNGF bind to p75NTR) pathway led to activation of apoptotic cascade and was significantly upregulated in the analysis. It is worth mentioning that studies showed that brain tissues affected by AD had both necrotic and apoptotic regions (47).

In Parkinson’s disease, the pathway enrichment analysis showed that the R-HSA-5632927 (defective mismatch repair associated with MSH3) pathway is one of the most significantly downregulated pathways with an FDR value of 0.002. Other downregulated mismatch repair pathways were diseases of mismatch repair, mismatch repair directed by MSH2:MSH3 (MutSbeta), mismatch repair, defective mismatch repair associated with MSH2, and mismatch repair directed by MSH2:MSH6 (MutSalpha). DNA damage was found to be one of the drivers of neurodegenerative diseases in which DNA repair mechanisms were involved (48).

One of the significantly enriched pathways in HD was R-HSA-2032785 (YAP1- and WWTR1 (TAZ)-stimulated gene expression), which had an FDR value of 0.00001. It is involved in the Hippo signaling pathway, in which the activation of the latter may affect neurodegeneration of the brain (49). The loss of *YAP1* may have the potential as a clinical marker for predicting neuroendocrine features and chemosensitivity (50). The R-HSA-6804759 (Regulation of TP53 activity through association with co-factors) pathway was upregulated with a significant FDR (0.004), where 13 proteins were involved. High levels of p53 protein encoded by TP53 were correlated with nerve cell apoptosis and neural damage (51). R-HSA-389977 pathway (post-chaperonin tubulin folding pathway) was also downregulated with an FDR value of 0.006. Other pathways, including R-HSA-351143 (agmatine biosynthesis), were reported for downregulation; it is known that agmatine binds to several target receptors in the brain and has been proposed as a novel neuromodulator (52).

### Biological pathways enriched between NDs: Neuroinflammation and cancer

Nine pathways were found common between AD and PD, 14 pathways were common between PD and HD, and six pathways were common between AD and HD only (Fig. 2, Table 2).

**Fig. 2.**
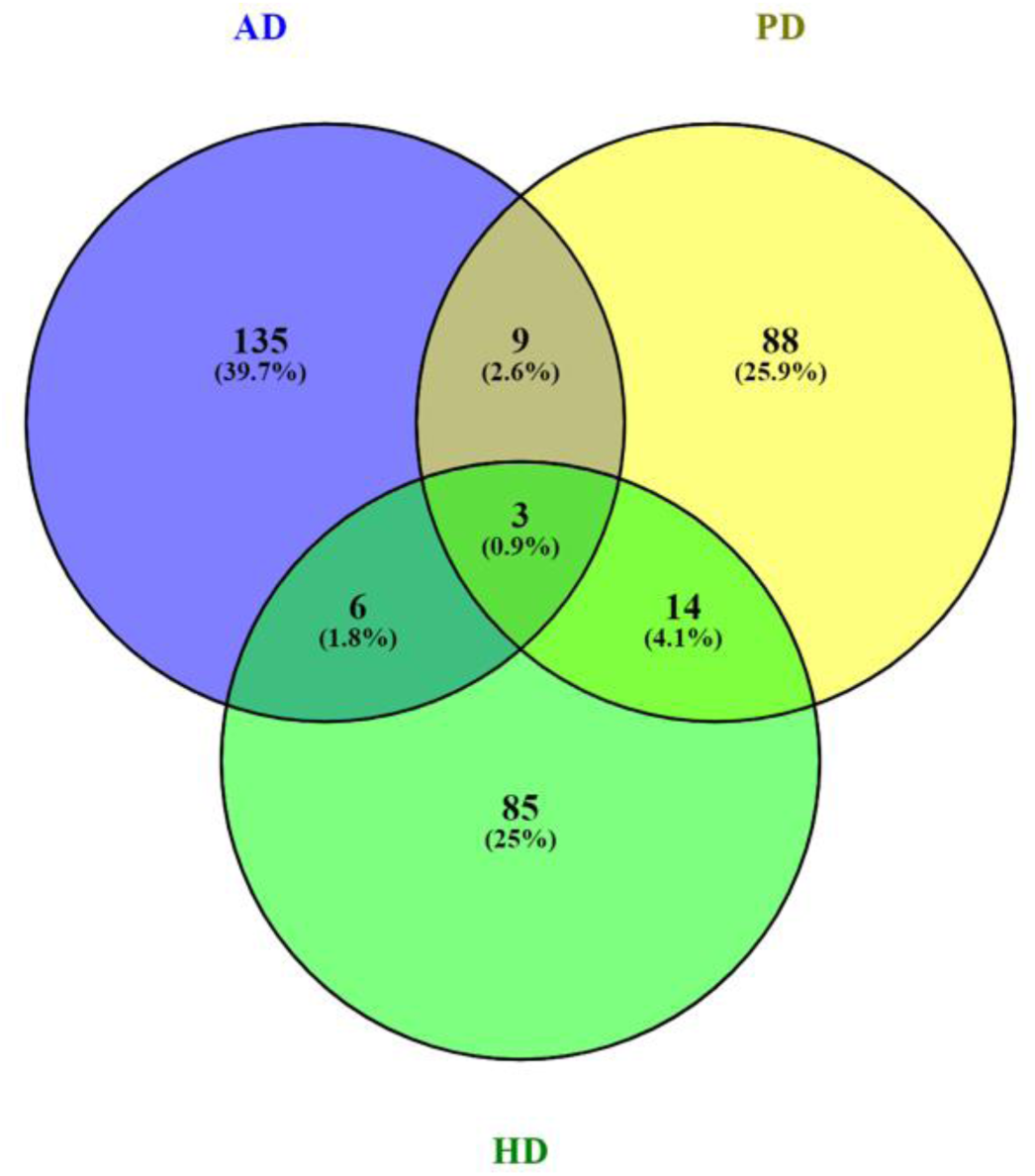
**Unique and common enriched pathways in AD, PD, and HD**

**Table 2:** List of mutual upregulated and downregulated pathways between each pair of NDs in the study.

|  | AD intersects PD | PD intersects HD | AD intersects HD |
| --- | --- | --- | --- |
| Upregulated Pathways | MyD88 cascade initiated on plasma membrane | Loss of Function of SMAD2/3 in Cancer | Circadian clock |
|  | MyD88 dependent cascade initiated on endosome | Loss of Function of *TGFBR1* in Cancer | Common pathway of fibrin clot formation |
|  | Regulation of localization of FOXO transcription factors | Metallothioneins bind metals | FOXO-mediated transcription of cell cycle genes |
| | Toll Like Receptor 10 (TLR10) Cascade | MyD88 deficiency (TLR5) | IκBα variant leads to EDA-ID |
|  | Toll Like Receptor 5 (TLR5) Cascade | MyD88:MAL (TIRAP) cascade initiated on plasma membrane | RUNX2 regulates bone development |
|  | Toll Like Receptor 7/8 (TLR7/8) Cascade | Response to metal ions |  |
| | TRAF6 mediated induction of NFκB and MAP kinases upon TLR7/8 or 9 activation | Signaling by TGF-β Receptor Complex in Cancer | |
|  |  | TNF signaling |  |
|  |  | Toll-like receptor 2 (TLR2) cascade |  |
|  |  | Toll-like receptor 4 (TLR4) cascade |  |
|  |  | Toll-like receptor TLR1:TLR2 cascade |  |
|  |  | Toll-like receptor TLR6:TLR2 cascade |  |
|  |  | Toll-like receptor cascades |  |
| Downregulated Pathways | Rap1 signaling | Activation of BMF and translocation to mitochondria | Defective GSS causes GSS deficiency |
|  | Regulation of pyruvate dehydrogenase (PDH) complex |  |  |

Seven of nine mutual pathways were upregulated between AD and PD, while two were downregulated. Most upregulated pathways were related to Toll-like receptors (TLRs) cascades. These included MyD88, TLR5, TLR10, and TLR7/8 cascades. Another upregulated pathway was the regulation of the localization of FOXO transcription factors. *FOXO* stands for forkhead box O, and they are transcription factors that influence various cellular functions such as intracellular signaling, metabolism, proteostasis, and cell cycle arrest (53). FOXO transcription factors were correlated with type 2 diabetes and neurodegeneration (54). The Ras-associated protein-1 (Rap1) signaling pathway was downregulated in AD and PD. Rap1 signaling pathway may enhance tumor invasiveness and metastasis in cancers such as breast cancer, pancreatic cancer, and leukemia (55).

The highest number of mutual pathways were observed between PD and HD. It may suggest more common pathophysiological features between the two diseases. Thirteen out of fourteen mutual pathways were upregulated. TLR2, TLR4, TLR1:TLR2, and TLR6:TLR2 cascades were significantly upregulated with FDR below 0.05. Tumor necrosis factor (TNF) signaling was another upregulated pathway in the study. TNF elevation was associated with inflammation and involvement in AD and PD (56). These findings suggest that it may also contribute to the pathophysiology of HD.

Five of six pathways were mutually upregulated between AD and HD. Remarkably, the fibrin clot formation pathway was upregulated and there has been evidence of the association of fibrin with neuroinflammation and neurodegenerative disease progression (57). FOXO-mediated transcription of cell cycle genes was upregulated in both diseases. It is worth mentioning that the *FOXO1* gene was upregulated in both PD and HD, *FOXO4* was upregulated in both AD and PD, while *FOXO6* was upregulated in PD only. The role of FOXO transcription factors in neurodegenerative diseases was thoroughly reviewed (58).

Three distinct pathways were shared between the three neurodegenerative diseases. They were found to be related to the immune system regulation and cancer (Supplementary Table S1.4, Additional File 1). They are TRAF6 mediated NF-kB activation, RUNX3 regulates BCL2L11 (BIM) transcription, and TAK1 activates NFkB by phosphorylation and activation of nuclear factor-κB (IκB) kinase (IKK) complex.

Most of the enriched pathways involve immune response regulation and inflammation, such as NF-κB transcription factor activation and signaling. In concordance with these findings, out of the three mutual pathways significantly enriched in the three diseases, two mutually enriched pathways are involved in inflammation, namely TRAF6-mediated and TAK1-mediated NF-κB activation pathways. We will detail the three pathways to propose genes involved in them as key mutual drug targets in the discussion section.

### Among thousands of unique differentially expressed genes, only 274 genes were common

In this study, DEGs were defined as those with an adjusted P-value less than 0.05. DEGs for each neurodegenerative disease in the study were reported and visualized, highlighting key mutual drug targets involved in the three enriched mutual pathways (Fig. 3). In AD, 943 genes were downregulated, and 861 genes were upregulated. Regarding PD, 3802 genes were downregulated, while 3539 genes were upregulated. For HD, 1938 genes were downregulated, and 2401 genes were upregulated. A detailed list of the DEGs in the three NDs can be found in the Supplementary Information Tables S2.1, S2.2, S2.3 (Additional File 2).

**Fig. 3.**
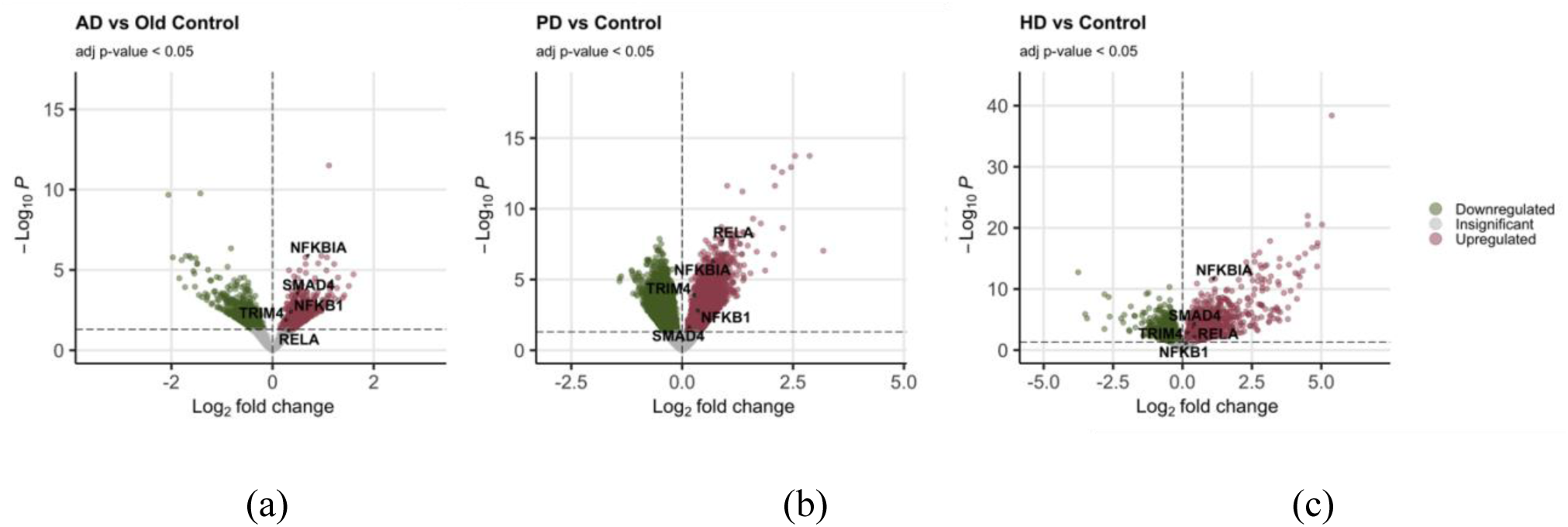
**Volcano plots for DEGs in (a) AD, (b) PD, (c) HD**

Overall, 274 mutual DEGs have been identified between the three NDs, as shown in Fig. 4. The mutual DEGs are reported with their P-adjusted and log fold change values from each disease dataset (Supplementary Table S2.4, Additional File 2). Mutual upregulated genes across the three diseases were 156, while mutual downregulated genes were 118. According to KEGG classification, most mutual DEGs fall under the following categories: enzymes, membrane trafficking proteins, transcription factors, and cell cycle proteins. For example, 58 of the DEGs were classified as enzymes, 22 as membrane trafficking proteins, and 18 proteins as transcription factors (Supplementary Table S2.5, Additional File 2).

**Fig. 4.**
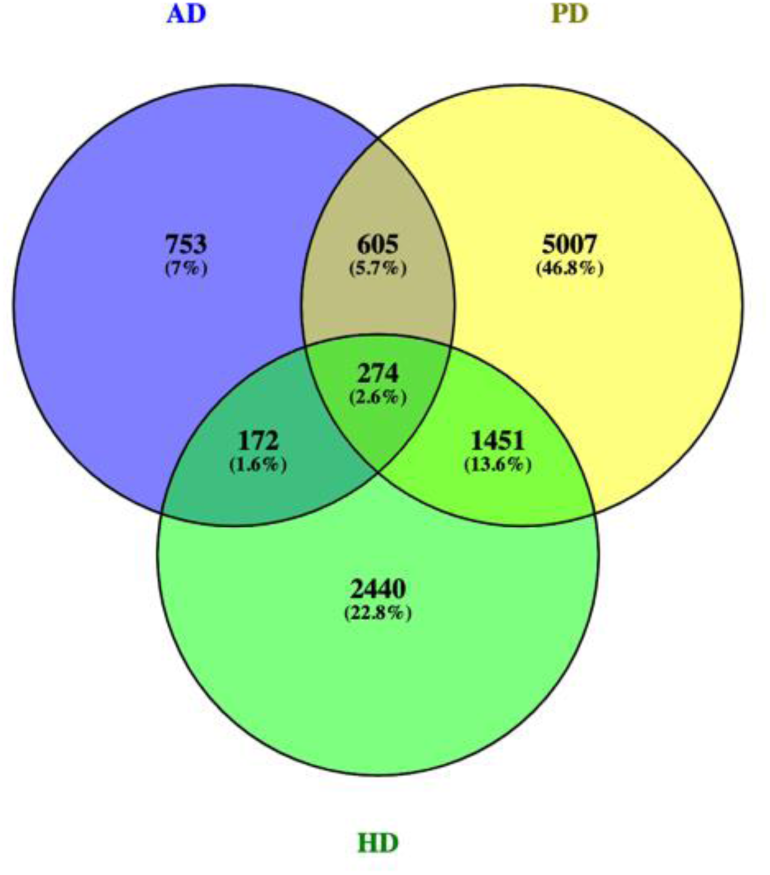
**Common DEGs between AD, HD, and PD**

### Key mutual drug targets across the three diseases

The genes involved in the three mutual pathways were screened (Supplementary Information Table S1.5, Additional File 1). To be considered a key mutual drug target, a gene had to be involved in a mutually enriched pathway and differentially expressed with the same pattern across the three diseases. The key mutual drug target genes found were *NFKB1*, *NFKBIA*, *RELA*, *TRIM4*, and *SMAD4*. The five genes were upregulated across the three diseases. The first four genes were involved in both pathways of NF-κB transcription factor activation and among the mutual DEGs between the three diseases. The pathway of RUNX3 regulating BCL2L11 (BIM) transcription is the third mutually significant enriched pathway in which the fifth gene, *SMAD4*, was a key mutual DEG involved in this pathway.

To test their potential as diagnostic biomarkers, the key mutual drug targets were explored by ROC curve analysis. Using the studied datasets, most genes had areas under the curve (AUC) of over 0.71, with the highest *NFKBIA* achieving an AUC of 0.93 in the Alzheimer’s dataset (Fig. 5). Using external datasets, most genes had an AUC of approximately 0.6 or more. ROC analysis using external validation datasets showed different AUC values for the selected genes (Fig. 6). This may be attributed to the differences in the sample size and brain regions between the datasets included in the primary analysis and the external validation datasets. However, they were selected for validation due to the similarities in the experimental design.

**Fig. 5.**
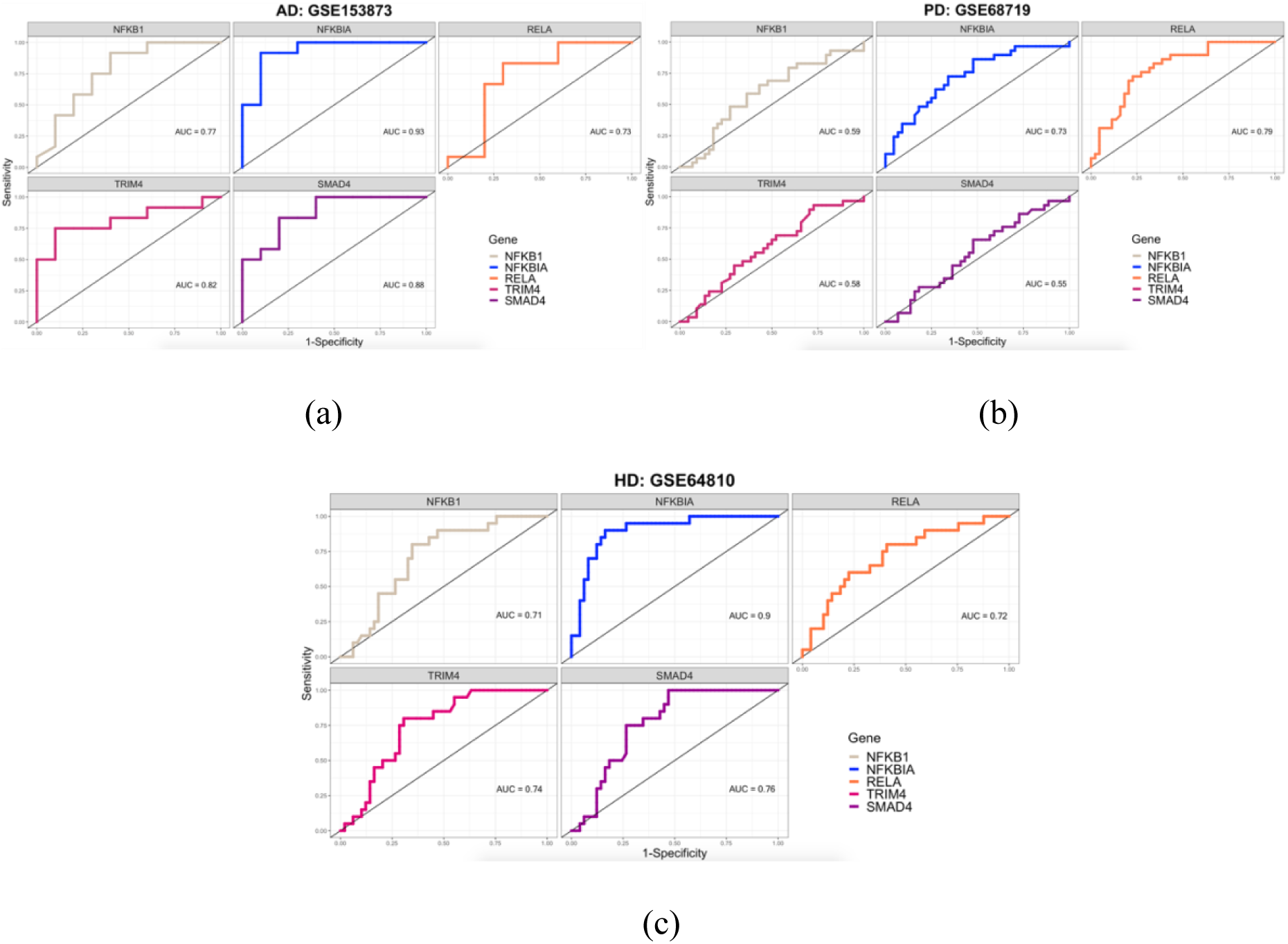
**Internal validation using ROC analysis curve for the selected DEGs in (a) AD, (b) PD, (c) HD**

**Fig. 6.**
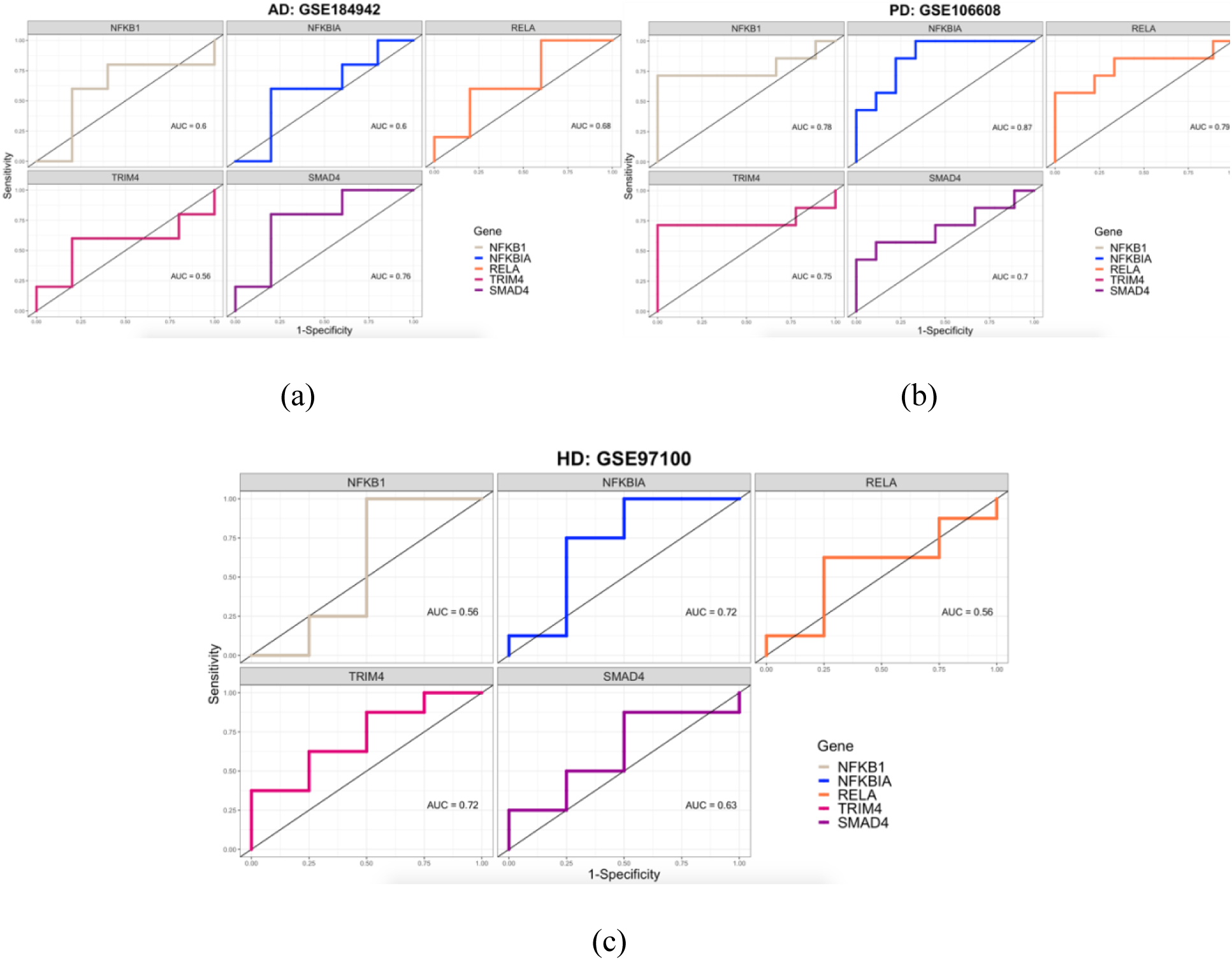
**External validation using ROC analysis curve for the selected DEGs in (a) AD, (b) PD, (c) HD**

### FDA-approved drugs repurposed for the treatment of the three diseases

According to the CMap dataset, only four genes of the five key mutual drug targets were found to have a measured gene expression in different perturbations. These were *NFKB1*, *NFKBIA*, *RELA*, and *SMAD4*. After filtering the CMap transcriptomic signatures for FDA-approved drugs, 73 drugs were found to downregulate these four key mutual drug targets (Supplementary Information Table S2.6, Additional File 2). After the exclusion of previously mentioned drug categories, filtered FDA-approved drugs included antihyperlipidemics, antihypertensives, antidiabetics, analgesics, diuretics, antiparkinson’s, and antipsychotics. The impact of each drug on the downregulation of any of the key mutual drug targets is represented by the intensity of the blue color in the figure below (Fig. 7). The darker the color, the more the drug downregulates the expression of the corresponding gene. One drug can affect the expression of more than one key mutual drug target.

**Fig. 7.**
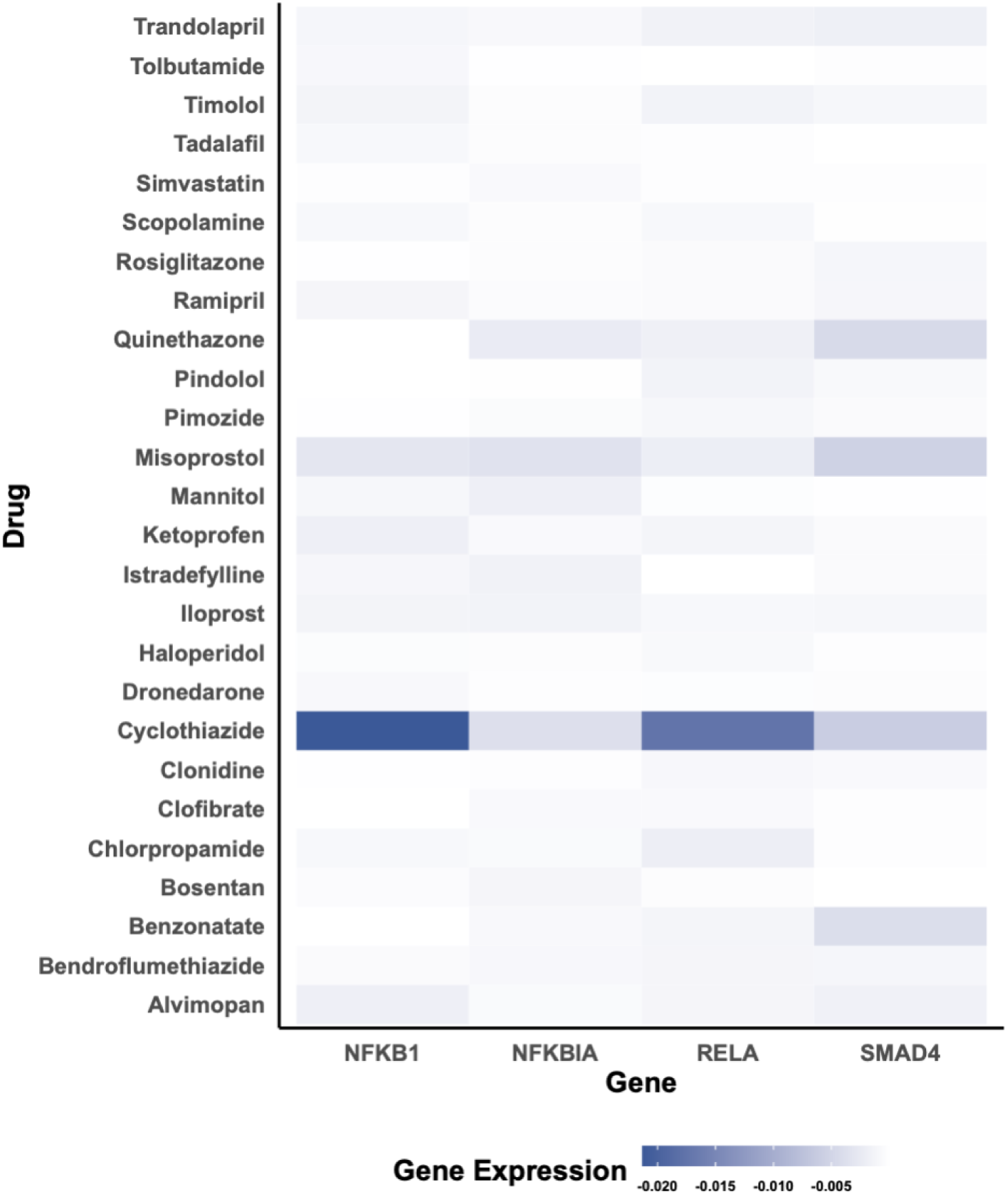
**CMap gene expression profile of key mutual drug targets to be downregulated by filtered FDA-approved in AD, PD, and HD.**

## Discussion

Due to their complex and progressive nature, research efforts are being expended to conclude the resemblances and discrepancies between different NDs. Early diagnostic tests and therapeutic interventions are sought to help limit the inevitable nerve damage threatening the patients’ quality of life and reduce the global burden of NDs. Most NDs share common pathophysiological pathways, such as inflammation. In addition to accumulating irregular peptides or proteins, neuroinflammation was verified to be involved in the progression of the three diseases under study (59–61). While local inflammation is one hypothesis, constant systematic inflammation with elevated proinflammatory and inflammatory mediators can also cause neuronal damage (62,63).

NF-κB is a B cell-specific inducible transcriptional factor that regulates the transcription of proinflammatory mediators such as chemokines and cytokines. Under normal conditions, NF-κB finetunes inflammation and maintains cellular protection. It is present in the central nervous system glial cells and cerebral blood vessels (64–66). NF-κB and its subunits reside inactivated by the inhibitor IκB in the cytoplasm. Canonical and noncanonical pathways activate the NF-κB members to be translocated to the nucleus and exert their regulatory actions (66). However, inducible activation of NF-κB was proved to be involved in neuroinflammation and tissue damage in Parkinson’s and Alzheimer’s diseases (64,67).

In our study, NF-κB activation was a notably significant pathway between the three diseases. *NFKB1*, *NFKBIA*, and *RELA*, a subunit of NF-κB, were significantly upregulated in the three diseases and *RELA* was involved in more than one pathway activating NF-κB. Furthermore, we found that two other mediators activate NF-κB transcription factors: TNF, TRAF6, and transforming growth factor β-activated kinase 1 (TAK1). Interestingly, the pathways of these two mediators are mutually upregulated in the three diseases. TRAFs and TNF receptor superfamily signal the activation of NF-κB. TRAFs have E3 ligase activity that is ubiquitin-dependent. Once TRAFs lysine at position 63 is ubiquitinated, they form a scaffold to signal downstream kinases (68). TAK1 and the nuclear IKK inhibitor are downstream kinases signaled by TRAF6 to activate NF-κB. TRAF6 was recently discovered to mediate both IL-1 and CD40 signaling (65). It plays a crucial role in the canonical activation of NF-κB so that other TRAF family members do not counteract its loss (68).

To further support the NF-κB activation, we found that the TNF signaling pathway is upregulated in Parkinson’s and Huntington’s diseases. TNF cytokines additionally induce the genes regulating inflammatory processes when the NF-κB pathway is activated (69). It is worth mentioning that in patients with Alzheimer’s disease, the NF-κB pathway might be activated in areas surrounding the β-amyloid plaques and stimulate the expression of TNF-α and IL-1ß cytokines. Activated NF-κB with the resultant neuroinflammation has been hypothesized as the leading cause of Alzheimer’s disease (64). Accordingly, inhibiting the NF-κB pathway may help prevent neuronal damage from the three diseases.

Toll-like receptors (TLR) are also involved in the signaling system of the NF-κB activation. Interleukin-1 receptor-associated kinase is phosphorylated and dissociated from myeloid differentiation factor 88 (MyD88)-dependent TLR signaling pathway. Phosphorylated interleukin-1 receptor-associated kinase consequently reacts with TRAF6 and promotes NF-κB pathway activation. TLR5 was one of our study’s upregulated genes of 274 mutual differentially expressed genes. TLR expression is enhanced in patients with Alzheimer’s disease and also in mouse models of Parkinson’s disease (59).

Based on the involvement of NF-κB activation in neurodegenerative diseases, several compounds, consistent to our findings, have been tested and proposed to limit or inhibit its activation, especially in AD. Non-steroidal anti-inflammatory drugs were one of the key agents to halt neuroinflammation, yet their adverse effects interfere with their action in preventing brain damage (61). Antioxidants and bioflavonoids are other alternatives to alleviate neuroinflammation and improve disease prognosis. Sulforaphane, resveratrol, loganin, and icariside II are naturally occurring compounds that showed promising results interfering with the NF-κB activation (61,63,70,71). In rats, the antihyperlipidemic atorvastatin was proven to repress the expression of NF-κB together with TLR4 and TRAF6 (72).

Research studies have highlighted an intriguing inverse relationship between the progression of neurodegenerative diseases and cancer in which having one disease may decrease the risk of the other (73,74). In the three neurodegenerative diseases under study, we found that the BCL2L11 (BIM) transcription pathway modulated by *RUNX3* is upregulated. BIM or BCL-2-like protein 11 (BCL2L11) belongs to the BCL-2 protein family that induces cellular apoptosis (75). *RUNX3* is a runt domain-containing transcription factor that regulates the transforming growth factor (TGF- ß) pathway and upregulates BIM expression. TGF- ß pathway induces apoptosis through *BIM* and *RUNX3* regulation. With activated *SMADs* and *FOXO3A*, RUNX3 enhances the transcription of BIM. *SMAD4* is a tumor suppressor gene that inhibits epithelial cell proliferation and is upregulated in AD, PD, and HD in our study. Hence, the TGF- ß pathway can be a tumor suppressor pathway in which *RUNX3* is considered a tumor suppressor gene, especially in gastric cancer (75,76). This can support the hypothesis of the relation between the three neurodegenerative diseases and cancer development. It should be noted that although *RUNX3* is involved in the development of proprioceptive dorsal root ganglion neurons in mouse models and expressed in cranial and dorsal root ganglia in human, the exact role of *RUNX3* in neuronal diseases is not fully understood yet (77,78). This may be correlated with aging as *RUNX3* expression is observed to be elevated with age (79). Runt-related transcription factors, *RUNX3* is expressed in multiple hematopoietic lineages as well as in numerous tissues, including cranial and dorsal root ganglia, thymus, chondrocytes, and the mesenchyme of epidermal appendages. *RUNX3* is required to properly develop cytotoxic T-lymphocytes and Langerhans cells (78).

The key mutual drug target gene set found comprised *NFKB1*, *NFKBIA*, *RELA*, *TRIM4*, and *SMAD4*. They were mutually upregulated and were involved in the mutually enriched pathways between the three diseases. The effect of FDA-approved drugs on the expression of *NFKB1*, *NFKBIA*, *RELA*, and *SMAD4* was evaluated by CMap, where one drug can affect the expression of more than one key mutual drug target. Conventional antihyperlipidemics, antidiabetics, analgesics, diuretics, antihypertensives, and antiparkinsons can be repurposed to treat the three studied neurodegenerative diseases. Some of these FDA-approved drugs were previously tested in one neurodegenerative disease. Thus, we propose testing their effectiveness against other neurodegenerative diseases. Simvastatin and clofibrate are two antihyperlipidemic drugs that were found to downregulate the expression of the four genes in the current study. Simvastatin can adjust the abnormal cholesterol metabolism associated with the three NDs (80). Clofibrate showed antioxidant and anti-inflammatory effects that can provide neuroprotection against degeneration (81). Rosiglitazone, tolbutamide, and chlorpropamide were three antidiabetic agents retrieved by CMap. Rosiglitazone prevented neuroinflammation and oxidative stress in animal models of AD (82). Thus, its neuroprotective potential can be tested on the other two NDs. Both tolbutamide and chlorpropamide are not currently used for diabetic patients but can be formulated into nanoparticles for their protective effect on AD (83). Another drug that can be repurposed and delivered via nano systems is the anti-inflammatory ketoprofen which previously showed treatment potential for AD and PD (84). Multiple diuretics and antihypertensives can be tested to treat NDs as per the results of the current study. These included thiazide diuretics: bendroflumethiazide, quinethazone, and cyclothiazide, osmotic diuretic mannitol, central alpha agonist clonidine, beta-blocker pindolol, angiotensin-converting enzyme inhibitors (ACEI): ramipril and trandolapril, and phosphodiesterase 5 inhibitor tadalafil. In a study on mouse models of AD, bendroflumethiazide was found to reverse AD-induced cognitive impairments and decrease amyloid beta plaques (85). Cyclothiazide strengthened AMPA receptors signaling which led to more suppression of amyloid beta of the interstitial fluid of AD mouse model (86). On a *Drosophila* model of PD, mannitol was observed to have neuroprotective effect and to decrease α-synuclein aggregations (87). Clonidine proved to enhance the clearance of mutant aggregates causing HD (88). Ramipril and trandolapril are two centrally acting ACEI that cross blood brain barrier that may target AD risk factors (89,90). Trandolapril may have improved cognitive functions in neurodegeneration (91). Tadalafil may play a role in decreasing neuroinflammation and enhancing neuroprotection (92). Calcium channel blockers such as dronedarone regulate calcium in the zebrafish model of AD (93). Iloprost is a prostaglandin vasodilator that helps protect pericytes and decrease demyelination. Thus, it can potentially slow down AD neurodegeneration (94). Misoprostol is a synthetic prostaglandin approved for treating gastric ulcer and was found to protect against neurotoxicity in a rat model of PD (95). Istradefylline is an approved adenosine A2A receptor antagonist for the treatment of PD during off-periods (96). Pimozide is an antipsychotic drug that proved to activate autophagy via a non-mTOR pathway, reducing tau aggregates in AD mouse models (97).

## Conclusion

Based on the current study, the significant pathways between AD, PD, and HD are inflammation-mediated pathways that activate the NF-κB transcription factor. Targeting the cycle of NF-κB activation may serve as a potential therapy for the three NDs simultaneously. The analysis also provideed further evidence for the decreased risk of patients with NDs having cancer. Both conclusions can be further supported by experimental validation by knocking out targets in these pathways and studying their impact in vivo. While in silico validation is another approach, the availability of data with a similar study design for each neurodegenerative disease and the suitable sample size was quite challenging, especially with autopsy samples from the brain tissue. Conventional drugs can be repurposed to modulate the expression of the mutual genes involved in the mutual pathways between the three diseases. Repurposed drugs, especially those already taken by the elderly for other indications, give hope to a large population of patients who suffer from progressive neurodegeneration and are unable to tolerate the unknown side effects of the newly discovered drugs.

## Author contributions

M.E. developed the pipeline, N.A. performed the biological analysis, E.B. conducted the project administration and conceptualization, M.E., N.A., and E.B. worked on the methodology, N.A. and M.E. worked on the original draft preparation, and E.B. conducted the manuscript review and editing.

## Author information

Center for Genomics, Helmy Institute for Medical Sciences, Zewail City of Science, Technology, and Innovation Nehal Adel Abdelsalam and Eman Badr Department of Psychiatry, University of Iowa, Iowa City, IA, USA

Muhammad Elsadany

Interdisciplinary Graduate Program in Genetics, University of Iowa, Iowa City, IA, USA

Muhammad Elsadany

University of Science and Technology, Zewail City of Science, Technology, and Innovation, Giza, Egypt

Eman Badr

Faculty of Computers and Artificial Intelligence, Cairo University, Giza, Egypt

Eman Badr

## Availability of data

All data generated or analyzed during this study are included in this published article and its supplementary information files. The datasets included in the analysis are publicly available in the GEO database under the following accession numbers: GSE153873, GSE68719, GSE64810, GSE70138, GSE184942, GSE106608, and GSE97100. The codes generated for the analysis are available at https://github.com/ComputationalBiologyLab/Shared-Pathways-and-Drug-Targets-in-Neurodegenerative-Diseases-Transcriptomic-Analysis

## Ethics Declaration

### Ethics approval and consent to participate

Not applicable

### Consent for publication

Not applicable

## Supplemental Files

Additional File 1

Additional File 1.xls

Table S1: Pathways enriched in AD, PD, and HD

Enriched pathways in three neurodegenerative diseases with FDR below 0.05. The file includes regulation of the enriched pathways (upregulated or downregulated), FDR, P values, and average fold change for Alzheimer’s, Parkinson’s, and Huntington’s diseases. It also includes mutual pathways between the three diseases and the genes involved in these pathways for each disease.

Additional File 2

Additional File 2.xls

Table S2: Differentially expressed genes in AD, PD, and HD

Differentially expressed genes in three neurodegenerative diseases. The file includes log fold change and adjusted P values for differentially expressed genes in Alzheimer’s, Parkinson’s, and Huntington’s diseases. It also has mutual differentially expressed genes between the three diseases. For the list of mutual DEGs, the classification of these genes is described according to the KEGG database. It includes full list of the FDA-approved drugs to downregulate key mutual targets retrieved from CMap with their drug category.

